# Probing behavior of the corn leafhopper *Dalbulus maidis* on susceptible and resistant maize hybrids

**DOI:** 10.1101/2021.10.21.465285

**Authors:** Pablo Daniel Carpane, María Inés Catalano

## Abstract

The corn leafhopper *Dalbulus maidis* is the main vector of the pathogens that cause corn stunt, a major disease of maize in the Americas. As host resistance is an efficient tool to control diseases, the findings of a previous report showed that some corn hybrids are resistant to *D. maidis*. In this work, we assessed the probing behavior of *D. maidis* on susceptible and resistant corn hybrids using EPG (Electrical Penetration Graph) technology. Fifteen-day-old females were monitored for 20 hours, with access to hybrids DK390, DK670, DK79-10, and DK72-10. Hybrids DK390 and DK72-10 showed resistance to *D. maidis* in phloem, since insects feeding on these hybrids presented more salivation events in phloem without subsequent ingestion, which are seen as failed attempts to ingest. A reduction of the total duration of phloem ingestion was observed, and accordingly of the time spent by insects with access to these hybrids on xylem ingestion. The hybrid DK390 also had mesophyll resistance, seen as less probing time and a higher number of probes of short duration. These findings support and are consistent with previous research, providing useful information to characterize maize hybrids resistant to *D. maidis*, and so to corn stunt.

## Introduction

Corn stunt is one of the most significant diseases affecting maize crop in the Americas due to its high potential to cause yield losses [1, 2, 3]. The increasing prevalence of corn stunt in the Americas [2, 4, 5] after its first detection [6, 7] is a major constraint for corn production. The mollicute *Spiroplasma kunkelii* Whitcomb is the pathogen mostly associated with corn stunt [3, 4, 8]. This pathogen is transmitted only by a few leafhopper species, being *Dalbulus maidis* (DeLong) the main vector species [8, 9, 10, 11] due to its distribution [8, 9, 11, 12] and transmission efficiency of *S. kunkelii* [13, 14].

One of the strategies to reduce yield losses resulting from corn stunt, as well as from other diseases, is the use of resistant genotypes [15, 16, 17, 18]. In pathosystems involving insect vectors, plant resistance can be aimed either to the pathogen or to the insect vector [19, 20], with the latter expressed as reducing preference (antixenosis) or survival (antibiosis). These resistance traits are usually present in other pathosystems [20, 21], and corn hybrids eliciting this behavior in *D. maidis* have been already found [22]. Both resistance traits reduce the duration of insect-plant interaction [19, 20, 21], which in turn might reduce the inoculation efficiency of the pathogen *S. kunkelii*, which increases up to 80% if such interaction is extended from 1 to 48 h [14].

However, the characterization of hybrids either triggering antibiosis or antixenosis fails to provide insights on the nature and location of plant resistance, hindering the development of breeding strategies like pyramiding resistance genes in search of durable resistance [18]. Since the transmission efficiency of *S. kunkelii* is directly related to the time spent by *D. maidis* probing in phloem [23], where *S. kunkelii* resides [24], a method capable of further characterizing the underlying features of plant resistance to *D. maidis* would ease the identification and characterization of the effect of resistance genes, thus contributing to designing even more effective strategies to control corn stunt.

The study of probing behavior using electrical penetration graph (EPG) technology recognizes the activities performed by insects while inserting their stylets into plant tissues, helping to identify the mechanism and location of the traits conferring plant resistance at the tissue level [25, 26]. This is particularly useful in pathosystems where the dynamics of pathogen transmission is highly related to the activities performed by insects on phloem [25, 27]. For instance, leafhoppers exposed to resistant genotypes typically spend less time probing from phloem than those probing on susceptible genotypes, decreasing the transmission efficiency of phloem inhabiting pathogens [21, 28]. Also, factors conferring plant resistance can be located in other tissues [26], and so they may be found by insects before reaching phloem [26, 29]. Hence, the study of probing behavior contributes to understanding the basis of plant resistance in pathosystems involving vectors and may assist plant breeders in selecting and characterizing resistant genotypes [20, 30, 31].

The objective of this work was to characterize the probing behavior of *D. maidis* adults with access to corn hybrids of different resistance to this insect species, in order to gain a better understanding of the mechanism and location of plant resistance traits in corn hybrids.

## Materials and Methods

A colony of healthy *D. maidis* was initiated from insects collected in Tucumán province, Argentina. This colony was reared on plants of the sweet corn variety “Maizón” at the CEBIO (BioResearch Center) of the UNNOBA-CICBA (National University of the North West of the Province of Buenos Aires—Scientific Research Commission of the Province of Buenos Aires), in Pergamino, Buenos Aires, Argentina. The colony was kept in aluminum-framed cages with a “*voile*” type nylon mesh, at a temperature of 25°C, under a photoperiod of 16:8 (light: darkness) hours [13]. Four maize hybrids were used in this study: DK670 and DK72-10 from the temperate region of Argentina (where corn stunt is not present), and DK79-10 and DKB390 from the tropical region of this country (where corn stunt is present). In a prior characterization of the resistance components of these hybrids to corn stunt [22], DK72-10 showed antibiosis and antixenosis to *D. maidis*, and DK390 showed antixenosis only. Moreover, DK79-10 was resistant to the pathogen *S. kunkelii*, and DK670 was considered susceptible to both *D. maidis* and *S. kunkelii*. The seeds used had no insecticides as part of the seed treatment. Plants were used at V3 (three leaves) stage.

Probing behavior of *D. maidis* was assessed using a Giga-8dd EPG-DC device (EPG Systems, Wageningen, The Netherlands) placed inside of a Faraday cage (made of aluminum mesh and wooden frame) to reduce the interference of external electrical noise. The Giga 8dd had eight probes, allowing to test eight insect-hybrid combinations (two per hybrid) per recording session. Ten to fourteen-day-old *D. maidis* females were anesthetized by single confinement in glass tubes (1.5 × 15 cm), and ice cooled for 30 minutes. The immobilized insects were then gently placed on a tethering stage with the help of a small brush. The dorsal side of the scutellum was tethered to a 12.5 μm diameter, 2-3 cm long gold wire using silver conductive paint for electric circuits (www.edelta.com.ar) under a binocular microscope. The other end of the gold wire was glued to a copper wire (around 0.5 mm diameter) soldered to a brass nail, assembling what is called the “insect electrode”. The length and diameter of the gold wire allowed free movement of insects, and some of them even attempted to fly away before being placed in contact with the plants. After tethering, insects were starved for 1 hour before the EPG recording session begun. Typically, 12-15 insect electrodes were mounted for each session, using only those moving steadily and so were not damaged during tethering. The “plant electrode” was a 2 mm diameter copper wire inserted into the moist soil of the pot that contained each plant. Plants were mounted by gently folding the newest expanded leaf around a 10 cm diameter cardboard cylinder and securing it with tape. The cylinder was then attached to a wooden stick inserted into the pot soil. This procedure exposed the abaxial side of the leaves to the insects and prevented the tissue exposed to the insect from moving during the EPG recording.

EPG recording sessions begun by mounting the “insect electrodes” in each of the eight “probes” of the EPG device and approaching the “plant electrodes” in such a way that insects and plants were two centimeters apart. The probe number assigned to each hybrid tested was randomized for each recording session. Once all insects and plants were in place, the EPG device was turned on and the pots were moved quickly (in less than 20 seconds) so the insects could reach the leaf tissue and start probing. The initial setup parameters were 10^9^ Ω input resistance and 50% gain, and the voltage supply and gain were adjusted independently on each probe for the first 30 minutes to ensure that the output voltage ranged from −5 to 5 V. In order to avoid differences in probing behavior that could be due to variations in the circadian cycle, recording sessions always started around 3:00 pm and lasted 20 h. Fifteen insect-plant replications per hybrid were recorded using new insects and plants in each replication.

EPG waveforms were acquired using Stylet+ software and stored in the hard drive of the computer. In this work, a “probe” was described as the insertion of insect stylets into plant tissues and seen in the EPG device as a deviation from a flat line at 0 V when insect stylets were not inserted. In most probes, insects would often perform more than one activity (each activity was seen as specific waveforms), each of them called “waveform events”. The observed waveforms (Figure 1 and Figure 2) matched closely those previously described for *D. maidis* [27], so the waveforms seen here were transformed into insect activities using that reference. The waveforms 1, 2, 3, 4, 5, and 6 in [27] were renamed C, G, id11, E1, E2, and id12, respectively, according to the usual nomenclature in DC-EPG systems [32, 33]. The sequence of waveforms was saved in ANA files in Stylet+, after visually assessing each waveform and saving their name and time of occurrence. Several parameters of probing behavior were extracted from ANA files using EPG-Calc 6.1.3 [34]. Additionally, the duration of the complete phloem salivation events before phloem ingestion (with the entire sequence shown in Figure 2B through 2F) was calculated, discarding short events of phloem salivation amidst phloem ingestion detected in some insects. Waveforms id11 and id12 were not further analyzed due to their unusual presence (less than 1% of the recording time).

**Figure 1:**
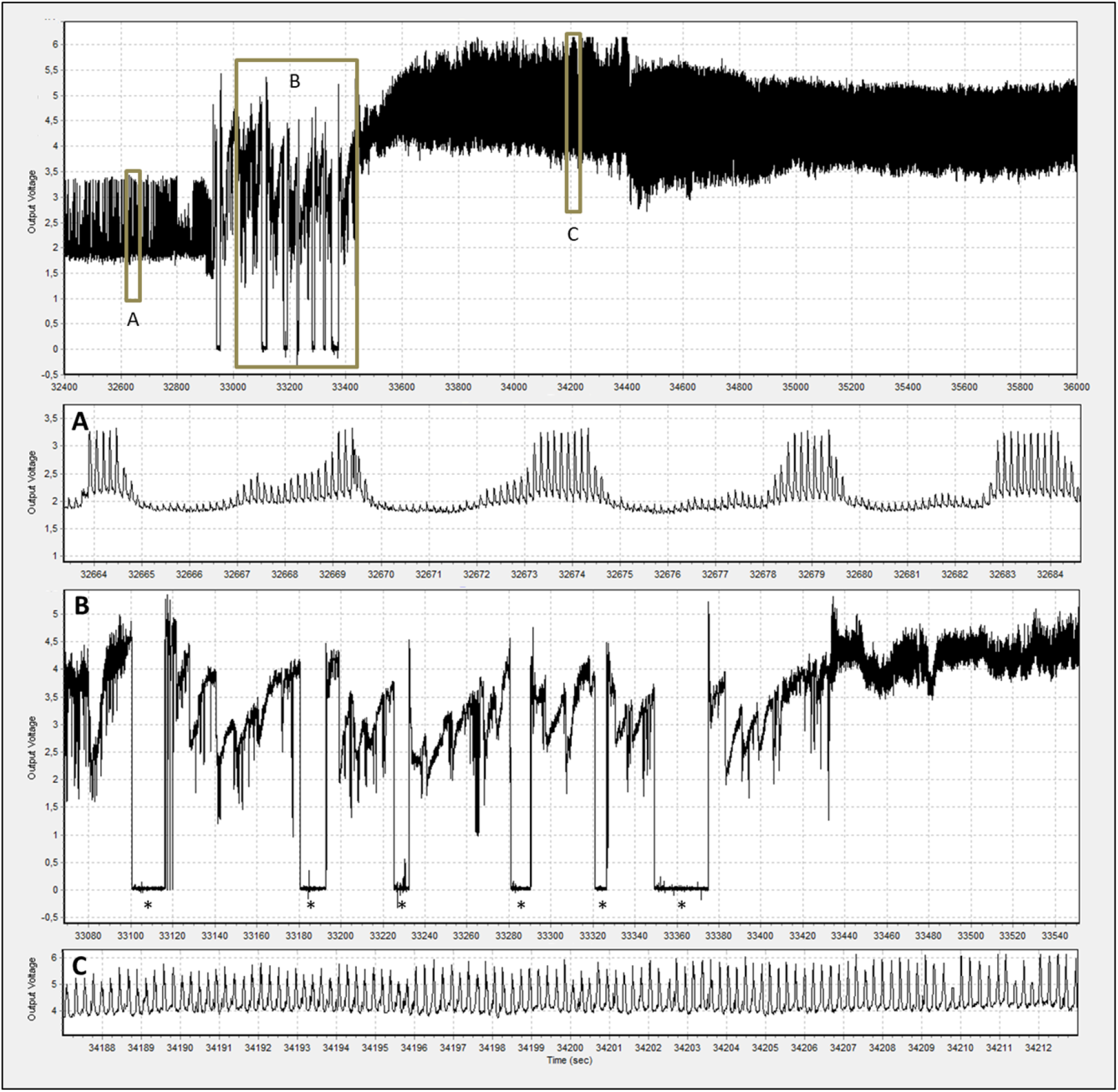
Samples of major waveforms displayed during probing of *D. maidis* females. A: phloem ingestion. B: a series of five short probes with pathway waveforms only, combined with brief non-probing periods (*), followed by a longer probe in which the insect begun to ingest from xylem (at around 33440 sec). C: xylem ingestion. the X axis indicates the time (seconds) from the beginning of the recording session.

**Figure 2:**
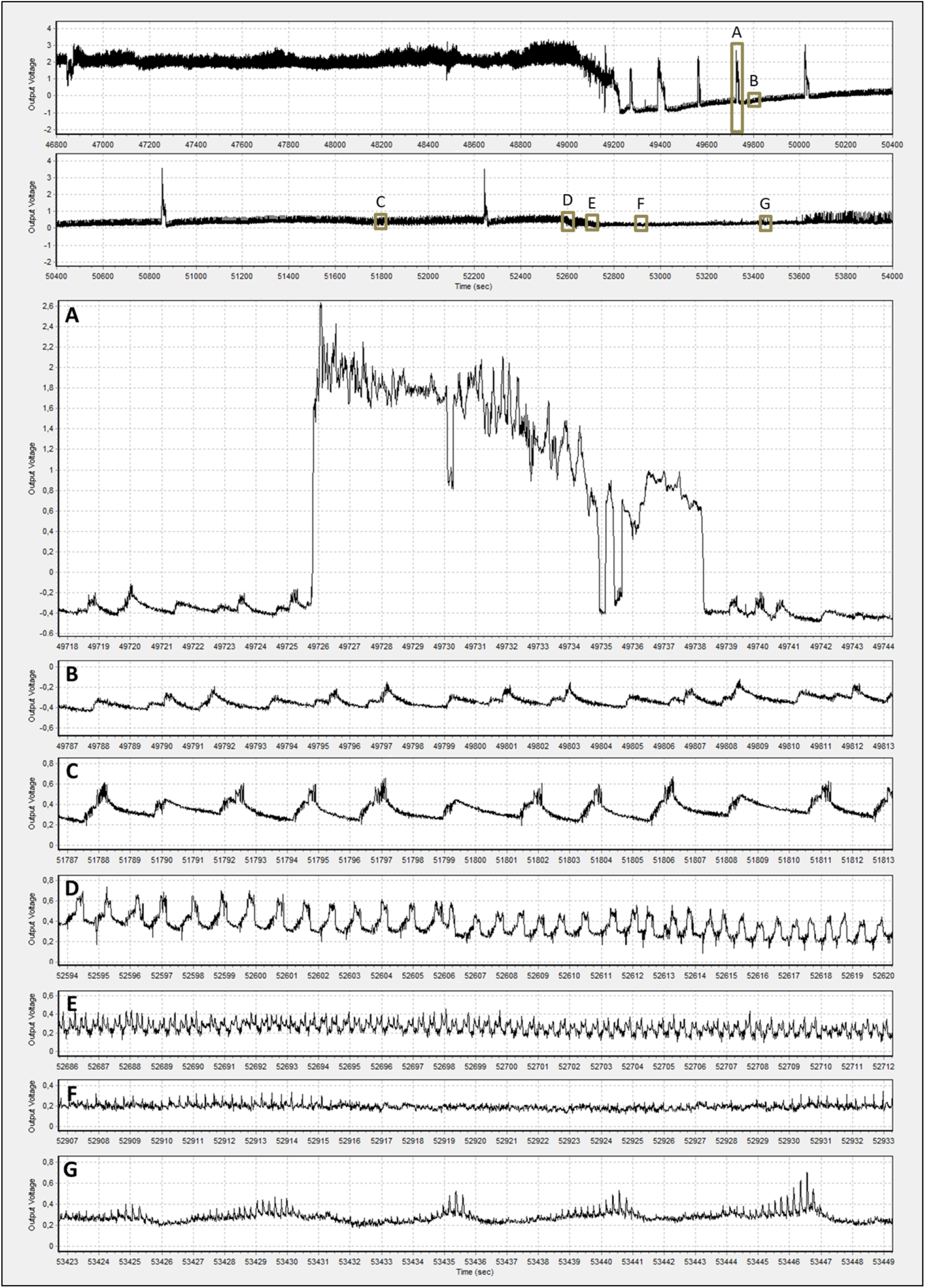
Progression of phloem salivation-ingestion of *D. maidis* insects with access to four maize hybrids. A: “spikes” of high amplitude and irregular shape, B-C: phloem salivation. D: end of phloem salivation. E: transition to phloem ingestion. F: early stages of phloem ingestion. G: phloem ingestion.

Statistical analyses were performed using Infostat [35]. The number of events of each waveform was analyzed using a Poisson distribution, and the remaining parameters were analyzed using a gamma distribution. In all the cases, the hybrid was the fixed effect, and the date of the recording session was the random effect. The models were adjusted using the nnet package [36] of the R language [37]. The significance of the differences in means was analyzed using Fisher LSD test.

## Results

The probing behavior of *D. maidis* was highly dynamic on the hybrids used, with the first probes occurring before the first minute, and a continuous succession of probes (mostly brief), with short non-probing periods, before insects begun ingestion from xylem. The number of probes (n_Pr) was highest in DK390 and lowest in DK72-10 (Figure 3A). The average duration of probing periods (a_Pr) was longest in DK670 and shortest in DK390 (Figure 3B), ranging from 12 to 36 min. The lowest total probing time (s_Pr) in DK390 was a result of shorter probes, despite the highest number of probes (Figure 3C). Insects feeding on this hybrid also had the lowest ratio of time to first phloem contact to the number of probes (a_tC>1E/Pr), implying that more probes were needed to contact phloem for the first time (Table 1). Finally, the time required to first contact phloem (T<1E) was lower in DK670 than in the other hybrids (Figure 3D).

**Table 1:** Parameters (Average +/- Standard Errors) of probing behavior of *D. maidis* insects with access to four maize hybrids. Values with the same letter are not significantly different according to contrasts in the mixed model test (α = 0.10).

| Variable | p value | DK670 |  |  | DK79-10 |  |  | DK72-10 |  |  | DK390 |  |  |
| --- | --- | --- | --- | --- | --- | --- | --- | --- | --- | --- | --- | --- | --- |
|  |  | Avg. | SE | Sig | Avg. | SE | Sig | Avg. | SE | Sig | Avg. | SE | Sig |
| a_NP | <0.0001 | 0.45 | 0.02 | A | 0.38 | 0.02 | B | 0.32 | 0.01 | B | 0.44 | 0.02 | A |
| m_NP | 0.1041 | 16.15 | 2.43 | A | 13.68 | 2.26 | AB | 11.21 | 1.75 | B | 16.19 | 2.50 | A |
| mn_NP | <0.0001 | 0.012 | 0.001 | D | 0.018 | 0.001 | B | 0.013 | 0.001 | C | 0.019 | 0.001 | A |
| mx_NP | 0.0017 | 3.75 | 0.94 | A | 1.70 | 0.43 | B | 1.62 | 0.41 | B | 4.52 | 1.15 | A |
| n_bPr | <0.0001 | 49.99 | 9.27 | B | 47.47 | 8.88 | B | 33.58 | 6.22 | C | 59.16 | 10.93 | A |
| m_Pr | <0.0001 | 2.57 | 1.01 | AB | 1.05 | 0.42 | C | 2.58 | 0.61 | A | 1.40 | 0.40 | BC |
| mn_Pr | 0.0815 | 0.065 | 0.032 | A | 0.029 | 0.014 | AB | 0.044 | 0.023 | A | 0.017 | 0.009 | B |
| mx_Pr | 0.0515 | 534.5 | 62.8 | A | 384.9 | 46.8 | B | 494.9 | 55.2 | A | 394.5 | 47.9 | B |
| n_C | <0.0001 | 87.7 | 10.6 | B | 88.0 | 10.9 | B | 72.9 | 8.9 | C | 113.1 | 13.8 | A |
| s_C | 0.0241 | 104.6 | 16.8 | B | 161.7 | 27.3 | A | 156.7 | 25.1 | A | 175.4 | 29.9 | A |
| s_G | 0.0945 | 108.8 | 17.4 | B | 114.1 | 29.2 | B | 173.3 | 41.9 | A | 211.9 | 55.3 | A |
| m_sgE1 | <0.0001 | 7.25 | 4.02 | B | 13.69 | 4.68 | A | 8.82 | 2.96 | AB | 7.36 | 5.02 | B |
| mn_sgE1 | <0.0001 | 3.44 | 2.01 | A | 4.63 | 1.78 | A | 1.92 | 0.62 | B | 1.39 | 0.00 | B |
| n_E1 | 0.0131 | 5.88 | 0.83 | B | 5.69 | 0.84 | B | 8.64 | 1.04 | A | 9.11 | 1.18 | A |
| a_E1 | 0.0514 | 33.87 | 4.86 | A | 38.84 | 5.55 | A | 30.81 | 3.99 | A | 23.63 | 3.44 | B |
| m_E1 | 0.0087 | 27.32 | 6.36 | A | 37.70 | 8.43 | A | 25.49 | 5.22 | A | 13.44 | 3.01 | B |
| n_E | 0.0518 | 9.11 | 1.03 | B | 8.89 | 1.08 | B | 12.22 | 1.22 | A | 12.07 | 1.34 | A |
| s_E | 0.0681 | 828.5 | 73.4 | A | 780.0 | 71.2 | A | 789.2 | 66.0 | A | 655.5 | 58.6 | B |
| a_tC>1E/Pr | 0.0993 | 139.5 | 25.3 | A | 135.4 | 25.2 | A | 117.2 | 21.7 | AB | 86.6 | 16.1 | B |

Description of events of probing behavior: a\_NP: average duration of non-probing, m\_NP: median duration of non-probing, mn\_NP: minimum duration of non-probing, mx\_NP: maximum duration of non-probing, n\_bPr: number of brief probes (shorter than 3 minutes), m\_Pr: median duration of probes, mn\_Pr: minimum duration of probes, mx\_Pr: maximum duration of probes, n\_C: number of pathway, s\_C: sum of pathway duration, s\_C: sum of xylem ingestion, m\_sgE1: median duration of single (not associated with phloem ingestion) phloem salivation, mn\_sgE1: minimum duration of single (not associated with phloem ingestion) phloem salivation, n\_E1: number of phloem salivation, a\_E1: average duration of phloem salivation, m\_E1: median duration of phloem salivation, n\_E: number of phloem, s\_E: sum of phloem duration, a\_tC>1E/Pr: ratio of time and number of probes required to first phloem contact.

**Figure 3:**
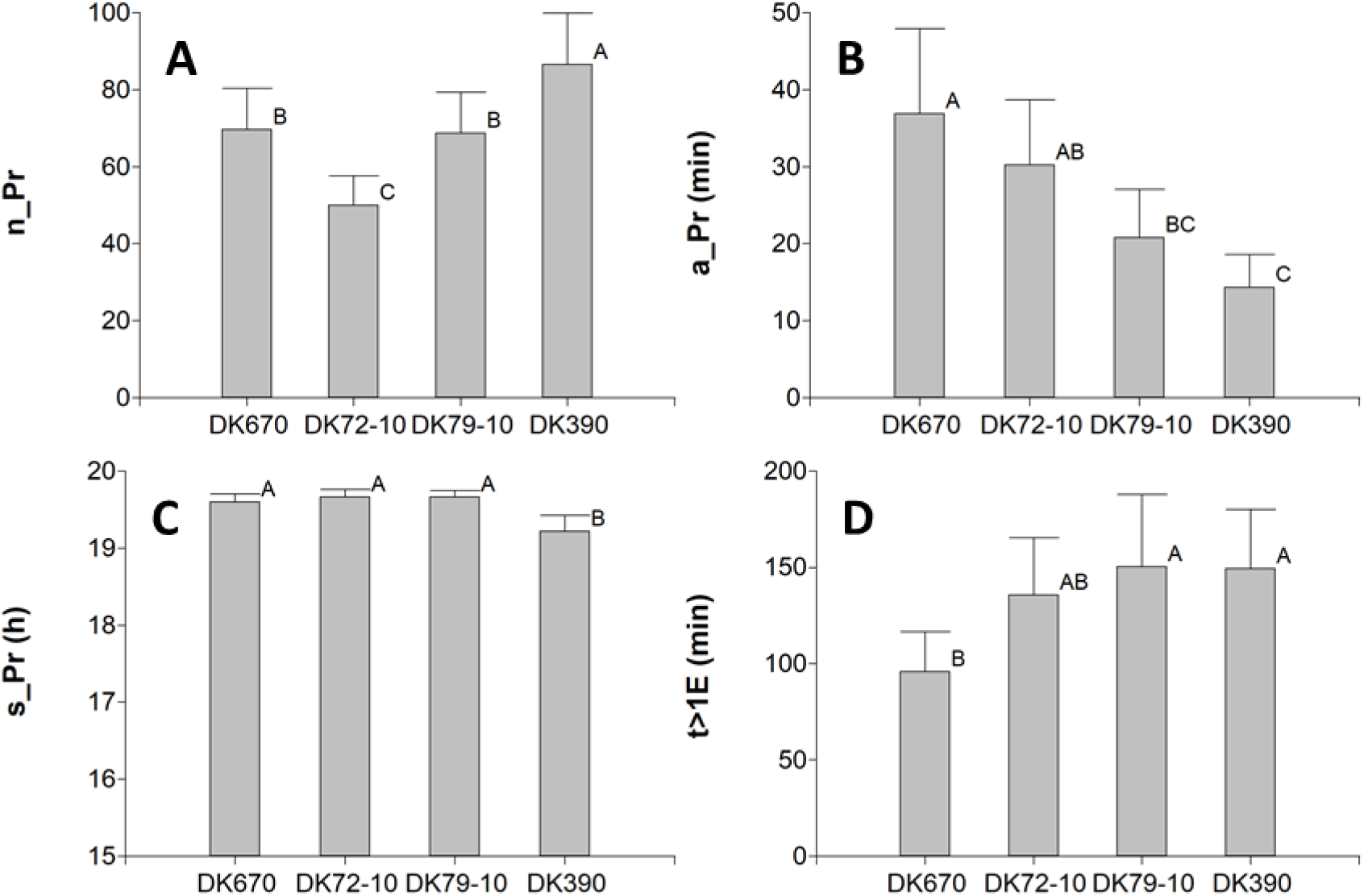
Parameters of probing behavior of *D. maidis* insects with access to four maize hybrids. A: number of probes (n_Pr), B: average duration of probes (a_Pr), C: total duration of probing (s_Pr), D: time to first phloem activity (t>1E). Values with the same letter are not significantly different according to contrasts in the mixed model test (α = 0.10). Bars indicate standard error of the mean.

In addition to the parameters described in Figure 3, other parameters related to the non-phloem phase (a_NP, m_NP, mn_NP, mx_NP, n_bPr, m_Pr, mn_Pr, mx_Pr, n_C, s_C) had the same pattern, with insects having more and shorter probes in DK390, and less time in probing and pathway in DK670 (Table 1).

The time spent on xylem ingestion ranged from 100 to 200 min, around 8 to 16% of the length of the recording session (1200 min). Xylem ingestion occurred within the first hour of probing and, for insects unable to ingest from phloem, within 5 hours. Half of the insects started xylem ingestion in less than 3 min, and by 10 min all insects had ingested from xylem. The only exception was one insect feeding on DK670, which started phloem salivation at 3.5 min in the first probe, and begun phloem ingestion at 58 min in the third probe that lasted 10.6 h. The total xylem ingestion time (s_G) was longest in DK390, intermediate in DK72-10, and lowest in the other hybrids (Table 1). Neither the number nor the average duration of xylem ingestion events was significantly different across hybrids (results not shown).

Phloem salivation took place later, after xylem ingestion. Half of the insects begun phloem salivation in less than two hours, with no major differences across hybrids (results not shown). All the insects had at least one phloem salivation event. The number of single salivation events (n_sgE1) was higher in DK390 and DK72-10 than in DK670 and DK79-10 (Figure 4A). Likewise, the average duration of single salivation events (a_sgE1) was shorter in DK390 and DK72-10 than in DK670 (Figure 4B), and the median and minimum duration of single salivation events have the same pattern (Table 1). Single salivation events are neither preceded nor followed by phloem ingestion, suggesting that they represent failed attempts to ingest from phloem. Conversely, there were no significant differences in the parameters of fraction phloem salivation events (frE1), which are indeed next to phloem ingestion events. Due to the lack of differences in the parameters of fraction salivation events, the parameters of total (single + fraction) salivation events responded similarly to single salivation events (Table 1). Additionally, insects with access to DK390 and DK72-10 required a higher number of probes to reach first phloem ingestion (n_Pr>1E2) compared to the other hybrids (Figure 4C), although the number of probes to first phloem contact (n_Pr>1E) was similar across hybrids (results not shown). The time to first phloem ingestion (t>1E2) was shorter in DK670 compared to the other hybrids (Figure 4D), directly related to the time to first phloem contact (t>1E) as shown before (Figure 2D).

**Figure 4:**
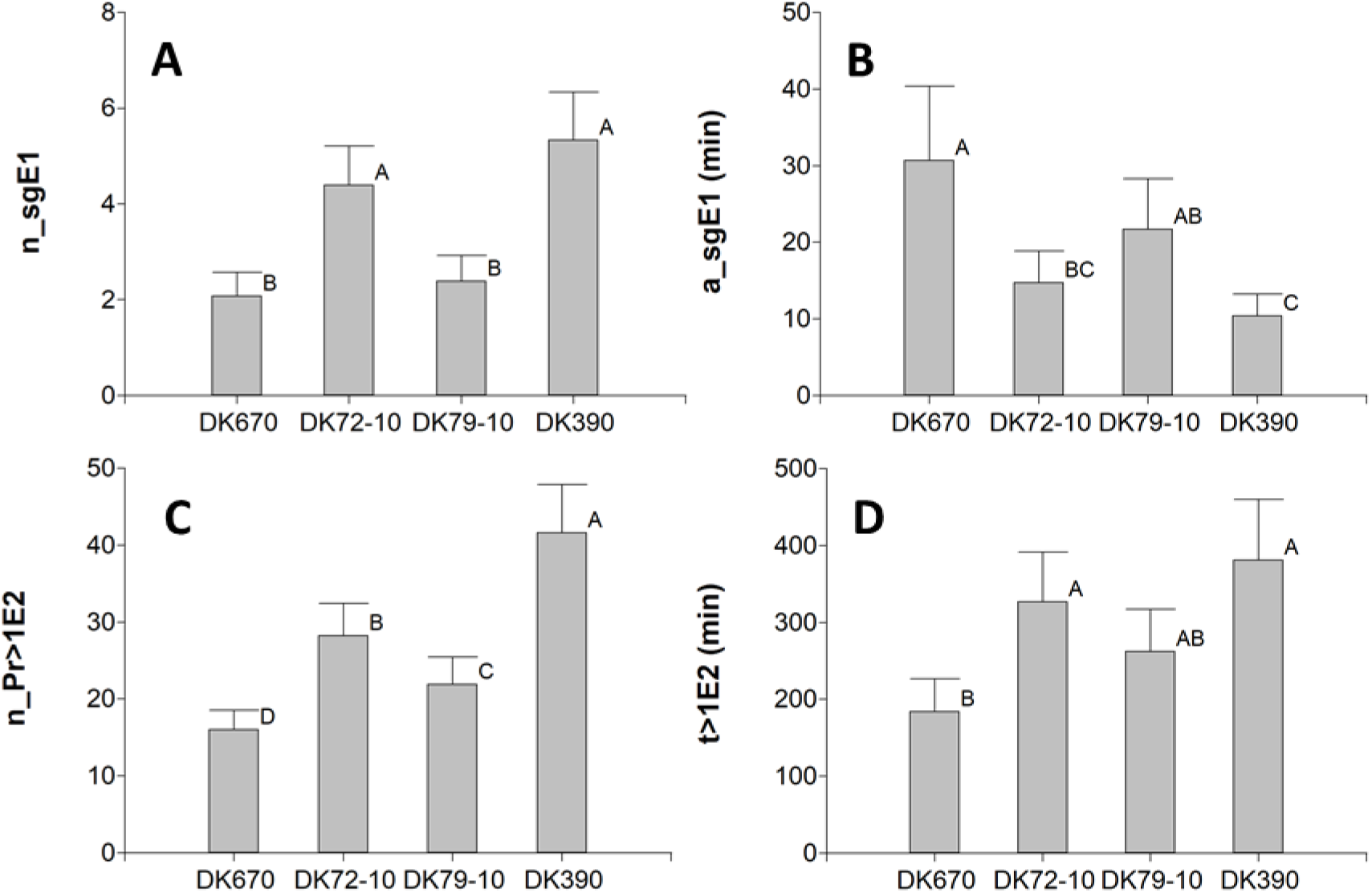
Parameters of probing behavior of *D. maidis* insects with access to four maize hybrids. A: average duration of single phloem salivation events, B: number of single phloem salivation events. C: number of probes before the first phloem ingestion. D: time to first phloem ingestion. Values with the same letter are not significantly different according to contrasts in the mixed model test (α = 0.10). Bars indicate standard error of the mean.

Phloem ingestion always followed phloem salivation, and it was the primary activity in the second half of the recording sessions. Half of the insects begun phloem ingestion in about four hours, with no major differences across hybrids (results not shown). There were usually two or three probes with phloem salivation before the first ingestion. All the insects in this study were able to reach phloem ingestion at least once. The time spent on sequences of phloem salivation + ingestion (s_E12) was shorter in DK390 compared to the other hybrids (Figure 5A). The total time spent on phloem (s_E), including the time spent on phloem salivation not followed by phloem ingestion, had the same pattern across hybrids (Table 1). The time spent on phloem ingestion (s_E2) was longest in DK670, followed by DK72-10, DK79-10, and shortest in DK390 (Figure 5B). Since most phloem ingestion events lasted more than 10 min, they were all considered “sustained”; therefore, the parameters related to this waveform (sE2) showed the same patterns across hybrids and were not further analyzed.

**Figure 5:**
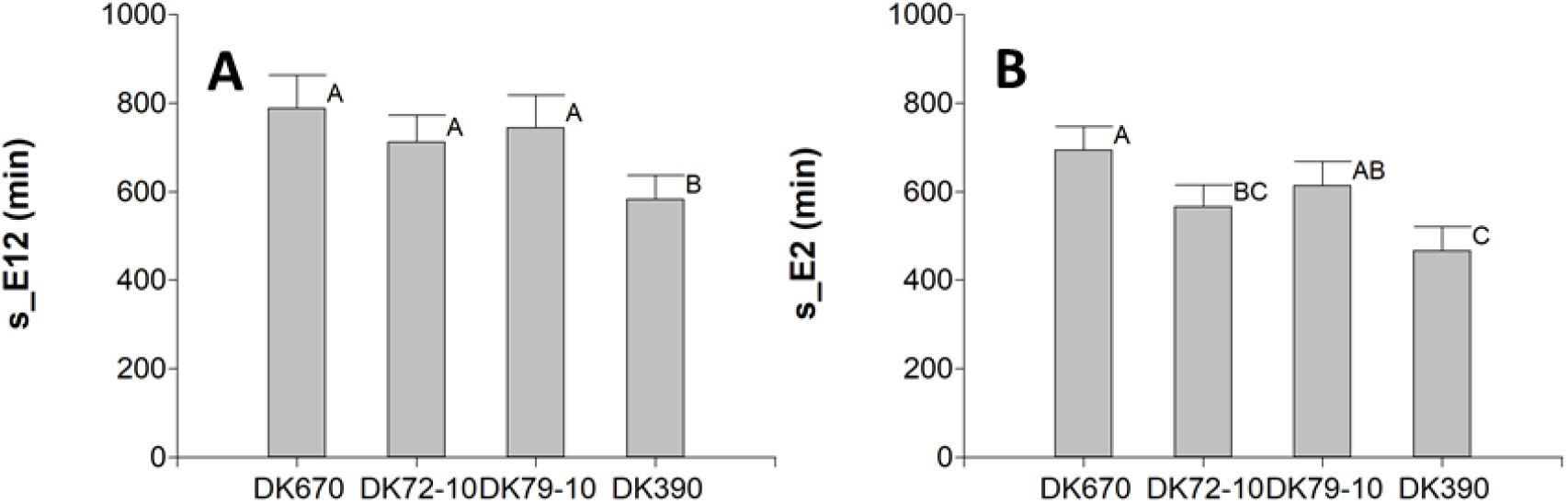
Parameters of probing behavior of *D. maidis* insects with access to four maize hybrids. A: sum of duration of phloem salivation + ingestion events, B: sum of duration of phloem ingestion events. Values with the same letter are not significantly different according to contrasts in the mixed model test (α = 0.10). Bars indicate standard error of the mean.

A summary of the parameters related to plant resistance to *D. maidis* for the hybrids used is shown in Table 2. Compared to the susceptible hybrid DK670, insects with access to DK79-10 showed an increase in the total time spent on pathway (s_C), and in the time to first phloem contact (t>1E). In addition to these changes, insects probing on DK72-10 increased the time spent on xylem ingestion (s_G), had more (n_sgE1) phloem salivation events of shorter duration (a_sgE1) not related to phloem ingestion, and required a higher number of probes (n-pr>1E2) and time (t>1E2) to reach phloem ingestion, all of this reducing the total time spent on phloem ingestion (s_E2). On top of the changes seen on DK72-10, insects probing in DK390 increased the number of brief probes (n_bPr), and the total number of probes (n_Pr), although the total time (s_Pr) and average probing duration (a_Pr) were reduced.

**Table 2:**
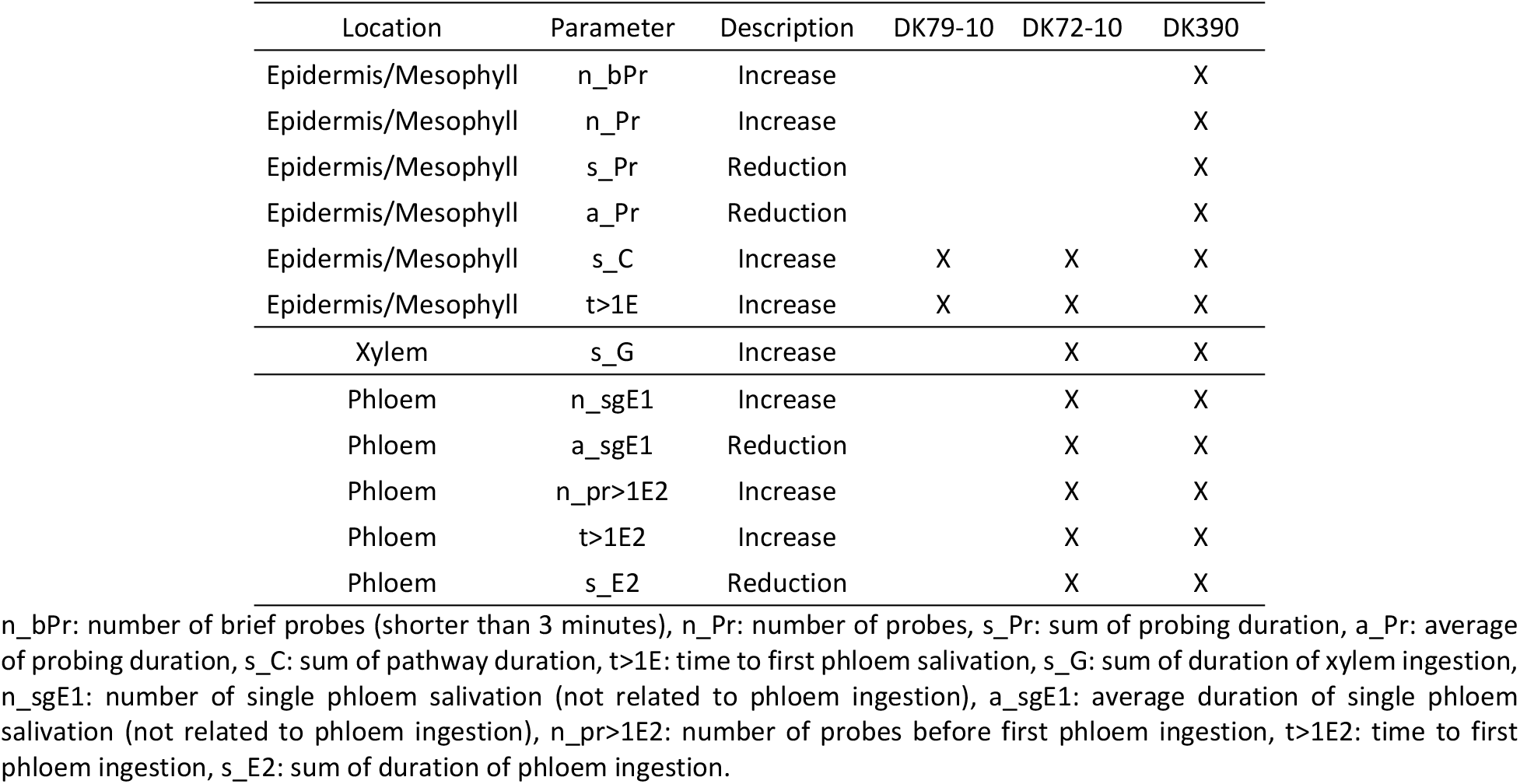
Parameters of probing behavior of *D. maidis* insects for hybrids showing significant differences from the susceptible hybrid DK670.

## Discussion

The reduction of total probing time due to shorter probes, despite an increase in the number of probes found here in DK390 for *D. maidis*, is similar to previous findings in the leafhoppers *Nephottetix virescens* [21, 30] in rice, and *Graminella nigrifrons* [38] and *Psammotettix alienus* [39] in several hosts, the planthoppers *Nilaparvata lugens* [31, 33, 40] in rice, and *Delphacodes kuscheli* [41] in corn and oats, and the aphids *Myzus persicae* [42] in oilseed rape, *Rophalosiphum padi* [29] in barley, *Acirthosiphon pisum* [43] in pulse species, and *Aphis glycines* [44] in soybeans. This behavior is observed as insects reject either non-hosts species or resistant genotypes before they contact phloem, implying that factors conferring plant resistance of the antixenosis type are located at the epidermis or mesophyll. Specific aspects of probing behavior related to resistance factors located at the epidermis, such as time to first probe and duration of first probe [29], were not found in this work with the hybrids tested, and insects probed quickly (within seconds) after placed in contact with plants, and several probes of short duration (less than one minute) took place in all genotypes during the very first period of insect-plant interaction. The lower time spent on pathway waveforms and time to first phloem contact by insects with access to DK670, compared to the other hybrids, coincide with the characterization of susceptible hybrids in other pathosystems, as in the leafhoppers *Nephottetix* spp. [28] in rice, the planthoppers *N. lugens* [31] in rice and *D. kuscheli* [41] in corn, and the aphids *R. padi* [29], A. *glycines* [44, 45] and *A. pisum* [43] in several host species, indicating an antixenosis-based resistance located in mesophyll found by insects before reaching phloem.

Xylem ingestion occurred in all the hybrids included in this work. As it has been already considered [25], this ingestion could be due to the need of insects to rehydrate after a dehydration period of about two hours during handling, before the start of the experiment. Furthermore, xylem ingestion at later stages of EPG recording sessions could be due to the need to rehydrate if the insects were not able to ingest from phloem, mainly in resistant hybrids, as it was seen mostly in DK390. This behavior was observed in *Nephottetix* spp. [28, 30] in rice, *G. nigrifrons* [38] in several hosts, *D. kusheli* [41] in corn, and *R. padi* [29] in several hosts. On the contrary, other work in *N. virescens* [21] and *A. glycines* [45] found no differences in the time spent on xylem ingestion, an aspect that could be related to either the insect species, the genotypes used, or the experiment setup as discussed above.

Insects with access to DK390 spent less time ingesting from phloem as compared to the other hybrids. This behavior of resistant genotypes or non-host species has been found in other species, such as the leafhoppers *G. nigrifrons* [38] and *N. virescens* [30, 21, 28], the planthoppers *D. kuscheli* [41] and *N. lugens* [40, 31], and in the aphids *A. glycines* [44, 45], *A. pisum* [43], *M. persicae* [42, 46] and *R. padi* [29]. Another aspect to consider regarding phloem ingestion is the proportion of insects reaching this stage. While all insects in our work reached phloem ingestion regardless of the hybrid, only a portion of the insects tested in most of the references cited reached phloem ingestion in leafhoppers [30], planthoppers [31, 33, 41], or aphids [43, 45]. This indicates that, in these cases, the factors conferring the antixenosis type of resistance are located in phloem and either prevent or reduce the ingestion of phloem sap.

The higher number and shorter duration of phloem salivation events not associated to phloem ingestion in DK390 and DK72-10 suggest the presence of a resistance factor in this tissue. However, the time spent on phloem salivation was not increased in these hybrids, as in earlier studies of the leafhoppers *N. virescens* [30] and *G. nigrifrons* [38], but disagreeing with the findings in the planthopper *N. lugens* [31] and the aphid *M. persicae* [46], which found an increase of the total time spent on salivation in resistant genotypes. Yet another aspect depicting a resistance factor located in phloem was seen as the first phloem ingestion was deferred in the resistant hybrids DK390 and DK72-10, similarly to the one described in the planthopper *D. kuscheli* [41] and the aphids *M. persicae* [42, 46], and *R. padi* [29], although the time to first phloem contact in the planthopper *N. lugens* [33] was similar both in resistant and susceptible rice varieties.

The proportion of insects reaching phloem is an aspect that allows to infer the efficacy of plant resistance traits to insect vectors in excluding the transmission of persistently transmitted pathogens. In this sense, while all the insects in this study reached this phase in all genotypes, some insects probing on resistant genotypes in *N. virescens* [30, 21, 28], *P. alienus* [39] and *N. lugens* [40, 31] did not reach phloem. This is clearly related to the insect species and plant genotypes used in each study, but perhaps also to the duration of the EPG recording session used to study the probing behavior of each insect. In this study, a 20 h recording session was selected as it is the timeframe for the efficiency of both inoculation and acquisition of the pathogen *S. kunkelii* by *D. maidis* to reach almost 100% [14], and prior research on this species [23, 27] indicated that insects actually needed this time to ingest from phloem, even in susceptible genotypes. In this sense, studies using short recording lengths would probably overlook the ability of vectors to feed from phloem, even in resistant genotypes, and to transmit pathogens accordingly. This aspect could be particularly useful in phloem-located pathogens and situations where the same genotype is deployed across large areas, and the plant hosts “chosen” by insects would be mostly other plants of the same genotype.

Compared to the susceptible hybrid DK670, DK79-10 showed some mesophyll resistance, seen as a longer time spent on pathway and a delay to contact phloem. This hybrid showed no antixenosis in a previous study [22], so it could be possible that either these parameters of probing behavior are a better alternative to detect traits conferring antixenosis, or this level of difficulty to access phloem plays no role in antixenosis seen as settling preference. The hybrid DK72-10 displayed mesophyll resistance like DK79-10, and phloem resistance seen as a higher number of phloem salivation events of short duration, not related to phloem ingestion. This implied a greater effort (both in number of probes and time) to reach phloem ingestion, resulting in less time spent on this activity and increasing xylem ingestion to avoid dehydration as seen in other species [28, 30, 45, 47]. Lastly, the hybrid DK390 altered *D. maidis* probing behavior in the same way as DK72-10, and reduced probing time due to shorter probes despite the increase in number, which relates to mesophyll-located resistance as discussed above.

The main current hypotheses of antibiosis and antixenosis to several insects are: a) the presence of toxins in either mesophyll or phloem tissues that are detected during insect probing and deter feeding [48, 49, 50, 51, 52], and b) the blockage of phloem sieve tube elements due to the inability of proteins in insect saliva to avoid clogging and maintain phloem salivation or ingestion [53, 54, 55, 56, 57]. Based on this, if the toxin is the underlying resistance mechanism, this would be present only in DK72-10, the only hybrid which reduced insect survival [22], as DK390 did not. Alternatively, a resistance mechanism based on a failure to avoid phloem clogging could be compatible with both hybrids. In addition, DK390 would also have a mesophyll-based resistance component of an unknown origin, since no reduced survival was seen in this hybrid [22].

While the characterization of the mechanisms of resistance to *D. maidis* in these hybrids is an open line for future research, another aspect to consider is how effectively they could prevent the transmission of the pathogen *S. kunkelii* by *D. maidis*. In this sense, the time to first phloem ingestion (50% at six hours) coincided in time with hybrids DK390 and DK72-10 being less preferred [22]. For this reason, although these hybrids would prevent phloem ingestion and would prompt the insects to flee in search of more suitable hosts, at least one event of phloem salivation has likely taken place, and so the pathogen *S. kunkelii* could be successfully inoculated [27]. Furthermore, insects moving to other plants of the same genotype would repeat this probing behavior, inoculating the pathogen to a high number of plants. An alternative would be that the inoculated pathogen is blocked by phloem proteins that promote phloem clogging, and so it would not be able to become systemic into the plant. The authors are not aware of any research in this direction at the present time, and hence this may also be an area of further research. Mesophyll-located resistance factors could also be a promising tool to prevent inoculation of *S. kunkelii*, since they operate at the first stages of insect-plant interaction, before insects make their first contact with phloem. This could potentially reduce the inoculation efficiency of *S. kunkelii* and can explain the slightly lower inoculation efficiency in DK390 (84%) compared to the other hybrids (100%) using five inoculative insects per plant [22]. This is also another potential area of research, which could benefit from the use of EPG monitoring with inoculative insects, and also help to dissect either *D. maidis* or *S. kunkelii* as the target of resistance in maize genotypes, as it has been discussed before [19, 20, 22].

This work provided a detailed characterization of maize resistance mechanisms to the corn leafhopper *D. maidis*. Sources of resistance were detected in mesophyll and phloem tissues, providing a tool to identify and combine sources of resistance, thus reducing the negative impact of corn stunt in field. Further research will attempt to identify genes related to this resistance, and to further characterize their impact in reducing the transmission efficiency of *S. kunkelii*.

## Acknowledgements

To Jazmín and Antonella Carpane, for their love, time, and patience while carrying out this work. To PIT-AP-BA (CIC) Res. 428/16 for providing funds for this project.

